# Particle-Only gECM Wafers Enable Cohesive, ECM-Rich Scaffolds Without Secondary Polymers

**DOI:** 10.64898/2026.06.20.733538

**Authors:** Shannon A. Blanco, Juliet O. Heye, Stephanie E. Schneider, Maxwell C. McCabe, Michael Floren, Corey P. Neu

**Affiliations:** Department of Mechanical Engineering, University of Colorado Boulder; Boulder, CO USA; Biomedical Engineering Program, University of Colorado Boulder; Boulder, CO USA; Mass Spectrometry Proteomics Shared Resource Facility, University of Colorado Anschutz; Aurora, CO USA; AlloSource; Centennial, CO USA; BioFrontiers Institute, University of Colorado Boulder; Boulder, CO USA

**Keywords:** Extracellular matrix (ECM), decellularized extracellular matrix (dECM), granular biomaterials, lyophilized scaffolds, tissue engineering

## Abstract

Granular extracellular matrix (gECM)-based biomaterials commonly contain polymer components to improve scaffold cohesion and handling during fabrication and use. However, these polymer hydrogel components may dilute ECM content and increase fabrication and regulatory complexity. This study evaluated whether particle-only gECM wafers could serve as a simplified alternative to hydrogel-based gECM scaffolds while maintaining structural, mechanical, and biological performance. Decellularized human cartilage and skin tissues were processed and fabricated into three scaffold formats: gECM hydrogels, freeze-dried gECM hydrogel wafers, and freeze-dried particle-only gECM wafers. Across fabrication methods, scaffold swelling, volume fraction, and stiffness were strongly influenced by both tissue type and fabrication approach. gECM hydrogels exhibited the greatest swelling and lowest stiffness, while gECM wafers displayed higher volume fractions and greater mechanical stiffness. Notably, gECM particle-only wafers achieved performance comparable to gECM hydrogel wafers despite the absence of a secondary polymer network. Particle-only wafers also maintained swelling behavior and structural properties over 3 months of dry storage at room temperature, with only modest decreases in stiffness. *In vitro* studies showed sustained cell viability over 14 days on particle-only wafers, with chondrocytes infiltrating cartilage wafers and fibroblasts remaining primarily surface-localized on skin wafers. In addition, particle-only wafers remained cohesive during implantation into a bovine cartilage defect model. These findings demonstrate that particle-only gECM wafers can achieve structural integrity, mechanical performance, and cytocompatibility without the need for an additional polymer network, highlighting a simplified and ECM-rich biomaterial platform. By eliminating polymer carriers and enabling dry storage with preserved function, this approach supports the development of off-the-shelf, translationally accessible gECM particle-only wafers for tissue engineering applications.

## 1. INTRODUCTION

Disease, injury, and trauma lead to tissue damage and degeneration, creating a clinical need for effective strategies to repair, replace, or regenerate damaged tissues. While allografts and transplants are commonly used for tissue replacement, their widespread application is limited by donor availability, cost, anatomical constraints, and restricted shelf life, which can delay timely access to suitable grafts [1,2]. Tissue engineering seeks to address these limitations by regenerating tissue rather than replacing it, through the development of biological substitutes composed of scaffold biomaterials and cells that restore, maintain, or improve tissue function [3]. Biomaterial scaffolds serve as a foundational component of tissue engineering systems, providing structural support while enabling cell migration, proliferation, and differentiation through mechanical and biochemical signaling [4,5].

Synthetic polymer scaffolds, including poly(lactic acid) (PLLA), poly(glycolic acid) (PGA), and poly(lactic-co-glycolic acid) (PLGA), enable precise control over mechanical properties and degradation rates [6]. However, these non-natural materials may elicit inflammatory responses or exhibit limited biocompatibility, potentially leading to cell death or impaired tissue integration. In contrast, non-synthetic biopolymers derived from biological components such as collagen and proteoglycans, support cell adhesion and signal cells to lay down their own extracellular matrix to replace the degrading scaffold. Hydrogels are among the most widely used scaffold types in regenerative tissue engineering due to their high water content and structural similarity to native soft tissues. These materials may be synthetic (PLLA, PGA, etc.) or non-synthetic (hyaluronan, gelatin, etc.) and typically rely on chemical or physical crosslinking to form three-dimensional networks [7].

Extracellular matrix (ECM)-derived biomaterials are commonly used to better recapitulate the native tissue environment, as the ECM provides essential structural, biochemical, and mechanical cues that regulate cell behavior. These materials are often processed into solubilized hydrogels; however, this approach can result in loss of native microstructure and insufficient mechanical strength [8–10]. To address these limitations, ECM hydrogels are frequently combined with synthetic or natural polymer components to form composite systems that improve stability and processability [11]. Despite these advantages, multicomponent systems introduce additional complexity. Polymer carriers may dilute ECM content, mask direct cell-matrix interactions, and require crosslinking strategies that increase fabrication complexity, and they may pose translational challenges related to manufacturing and regulatory approval [4,12].

An alternative approach is the development of non-synthetic, non-polymer ECM scaffolds, in which structural cohesion is achieved using ECM particles alone. By eliminating the hydrogel component, particle-only ECM scaffolds retain a higher density of biologically relevant ECM components such as collagen and proteoglycans, while simplifying fabrication by avoiding chemical modification and crosslinking. Particle-based systems still provide control over porosity, scaffold size, and geometry, and allow for flexible use of donor tissue sources. Prior work in the Neu laboratory has demonstrated the feasibility of incorporating ECM particles into composite hydrogel systems [13]; however, the ability of ECM particles to form cohesive scaffolds independently remains largely unexplored. Microgel-based granular hydrogel systems provide precedent for particle-based materials, demonstrating that densely packed microgels can form cohesive, load-bearing structures through physical interactions [14,15]. Similarly, processing approaches such as freeze-drying (lyophilization) have been widely used to generate porous, structurally stable biomaterials and may offer a route to stabilizing ECM particle assemblies while enabling dry storage [16–18]. These concepts suggest that ECM particles may form cohesive scaffolds through dense packing and contact-mediated interactions, even in the absence of a secondary polymer network.

The primary objective of this study is to biofabricate, characterize, and validate particle-only gECM wafers for use in both *in vitro* and *in vivo* tissue engineering applications. We focus on cartilage and skin as two compositionally and functionally distinct tissues that serve as test cases for this platform. Cartilage is a highly specialized, load-bearing tissue dominated by a dense collagen II and proteoglycan-rich matrix, supporting mechanical integrity and resistance to deformation. In contrast, skin is a more compliant, fibrous tissue composed primarily of collagen I, with structural organization that supports flexibility and remodeling. Together, these tissues represent contrasting extremes in ECM composition, structure, and mechanical function, enabling evaluation of whether particle-only scaffolds can be generalized across diverse tissue types. We first examine properties of decellularized ECM granules and architecture of particle-only wafers to establish a foundation for cohesion. Following this, we evaluate the mechanical and structural characteristics of particle-only wafers, using hydrogel-based constructs fabricated from the same tissue source as a benchmark for comparison. We hypothesize that particle-only gECM wafers fabricated via freeze-drying can achieve sufficient structural and mechanical performance to function as standalone biomaterials, while offering a simplified and potentially more translationally favorable platform. To assess translational stability, a shelf-life study was conducted in which cartilage and skin particle-only gECM wafers were evaluated over a 3-month timeline to determine whether structural and mechanical properties were preserved over time. Additionally, we evaluated cytocompatibility and an initial joint implantation model to assess biological performance and handling in a relevant setting. By demonstrating functionality across multiple tissue types, this work aims to establish particle-only gECM wafers as a viable foundation for future ECM-based biofabrication strategies and translational development.

## 2. METHODS

### 2.1 Decellularization and processing

Human cartilage and skin tissues were obtained from AlloSource. Cartilage samples were collected from 6 donors (5 male, 1 female; ages 37, 37, 38, 38, 39, and 37), and skin samples were collected from 5 donors (4 male, 1 female; ages 40, 67, 68, 74, and 67). Skin tissue was priorly decellularized at the AlloSource facility. Cartilage samples were decellularized in-house using a viral inactivation protocol [19]. Cartilage was cut into ∼4 mm^3^ pieces and agitated in 0.2M hydrochloric acid and 1.0M sodium hydroxide solutions for 2 hours each, followed by water and PBS rinsing until tissue demonstrated neutral pH. Following decellularization, cartilage and skin tissues were lyophilized and pulverized (Tissue Lyser III, Qiagen), then size-sorted using stainless steel sieves to obtain particles <250 µm in diameter. Decellularization was quantified using a PicoGreen double-stranded DNA Assay (Thermo Fisher Invitrogen). Successful decellularization was defined as residual DNA content below 50 ng per mg of dry tissue [20].

### 2.2 Particle-only gECM wafer formation

Decellularized cartilage and skin particles were suspended in 1X phosphate-buffered saline (PBS) at a concentration of 30% w/v (weight/volume) particles to PBS and mixed until homogenous. Particle suspensions were packed into cylindrical PDMS molds (5 mm diameter x 1.5 mm height). Filled molds were frozen at -20°C for approximately 4 hours and subsequently lyophilized overnight (∼12 hours). Following lyophilization, wafers were carefully removed from the molds and stored at room temperature under dry conditions until further use.

### 2.3 gECM hydrogel scaffold formation

The gECM hydrogel formulations consisted of 10 mg/mL tHA packed with 0.2 g/mL lyophilized cartilage or skin particles. One syringe containing ECM particles in a neutralization buffer (DPBS + 1M NaOH) was mixed using a luer lock connector to a second syringe containing an equal volume of 20 mg/mL tHA. The resulting gECM hydrogel was packed into cylindrical PDMS molds (5 mm diameter x 1.5 mm height). A glass slide was placed on top of the mold and constructs were incubated at 37°C for 45 minutes to polymerize. Resulting scaffolds were carefully removed from molds and stored at 4°C in PBS.

### 2.4 gECM hydrogel wafer formation

Wafers were formed using the same method as the gECM hydrogel scaffold, but after crosslinking at 37°C for 45 minutes, glass slides were removed from the top of the PDMS mold and constructs were placed in the -20°C freezer for 4 hours and subsequently lyophilized overnight (∼12 hours). Following lyophilization, wafers were carefully removed from the molds and stored at room temperature under dry conditions until further use.

### 2.5 Proteomics

Lyophilized, decellularized cartilage and skin samples (0.5–2 mg) were homogenized in hydroxylamine (HA) buffer (1 M NH□OH–HCl, 4.5 M guanidine HCl, 0.2 M K□CO□, pH 9.0) and incubated at 45°C with shaking for 4 hours to extract ECM-associated proteins. Following centrifugation (18,000 × g, 15 min), supernatants were collected and stored at −80°C.

Protein extracts were digested with trypsin (1:100 enzyme-to-protein ratio) using a filter-aided sample preparation (FASP) protocol and desalted using Evotips. Peptides (∼200 ng) were separated using an Evosep One liquid chromatography system and analyzed on a timsTOF Pro mass spectrometer (Bruker) operating in PASEF mode.

Raw data were processed using MSFragger (FragPipe v21.1) with a Sus scrofa UniProt database and common contaminants. Carbamidomethylation (C) was set as a fixed modification, while oxidation (M, P), deamidation (NQ), pyro-glutamate formation (N-term Q), and N-terminal acetylation were included as variable modifications. Label-free quantification was performed using IonQuant with match-between-runs enabled, and results were filtered to a 1% false discovery rate at the peptide and protein levels.

Identified proteins were classified into cellular and extracellular compartments using Gene Ontology annotations and MatrisomeDB. ECM proteins were further grouped into collagens, proteoglycans, glycoproteins, and associated categories. Protein intensities were normalized to total signal per sample and log□□ -transformed prior to analysis.

### 2.6 Particle size and swelling

Cartilage and skin particle size distributions were measured in a laser diffraction particle size analyzer (Malvern Mastersizer 3000). Particles were suspended in either 100% ethanol or deionized water to assess non-swelled and swelled particle states, respectively. Measurements were obtained using red (632 nm) and blue (470 nm) lasers passed through the suspension. Volume-based particle size distributions were calculated using Mie scattering theory, assuming spherical particle geometry. Tissue specific refractive indices were 1.40 (cartilage [21]) and 1.41 (skin [22]), while the absorptive index (0.1 [23,24]) and density (1.00 g/cm^3^) were held constant across tissues. Results are reported as the volume percentage of particles within each size range.

### 2.7 Scaffold microstructure

Scanning electron microscopy (SEM) was used to characterize scaffold microstructure and pore morphology in both dry and hydrated states. For dry imaging, lyophilized wafers were mounted and sputter-coated with a 10 nm platinum layer (Leica EM ACE600) prior to imaging.

For hydrated imaging, wafers were first equilibrated in PBS for 48 hours, followed by graded ethanol dehydration (increasing ethanol concentrations) to minimize structural collapse [25]. Wafers were placed in 20% ethanol for 2 hours, 40% ethanol for 2 hours, 60% ethanol overnight, followed by 2 hours in 80% ethanol and 100% ethanol overnight. Samples were then critically point dried (Leica EM CPD300) to preserve native architecture before sputter coating and imaging [25].

SEM images (Hitachi High-Tech SU3500) were acquired at multiple magnifications to assess pore size, particle packing, and interparticle connectivity. Representative regions were selected from the scaffold surface and cross-section to evaluate structural heterogeneity.

### 2.8 Scaffold topology

Surface topology and roughness of scaffolds were quantified using a laser scanning microscope (Keyence VK-X1000 Series). Both dry and hydrated scaffolds (hydrated in PBS for 48 hours) were analyzed. Three-dimensional surface profiles were acquired over three regions per wafer to account for spatial variability. Average surface roughness (Sa) was extracted from the 3D surface reconstructions using the Keyence analysis software. For hydrated samples, excess surface fluid was carefully removed prior to imaging to minimize optical artifacts. All measurements were performed using 20X magnification and scan settings were kept constant across samples to enable direct comparison.

### 2.9 Volume fraction

Cartilage and skin scaffolds were hydrated in PBS for 48 hours prior to imaging. Scaffolds were stained with Ghost Dye 710 (Cytek Biosciences) for 30 minutes and imaged using confocal microscopy at (Nikon A1R, 20x objective, NA=0.75). Z-stack images were collected with 5-micron slice spacing at 3 different locations per sample. Image analysis was performed using a custom MATLAB algorithm. Area fraction was quantified over a 15 µm depth using the highest-intensity slice and the two subsequent slices.

### 2.10 Swelling

Cartilage and skin scaffolds were submerged in PBS for 1 week. For the gECM hydrogel wafer and particle-only gECM wafer, scaffold height, diameter, and mass were measured in the dry state and after hydration at 30 minutes, 1 hour, 8 hours, 24 hours, 48 hours, and 1 week to assess time-dependent swelling behavior. For the gECM hydrogel, measurements were taken at t0 (immediately after fabrication), 1 hour, 24 hours, 48 hours, 1 week, and after lyophilization (dry). Surface area was measured from thresholded (Image J) stereoscopic images (Leica).

### 2.11 Compressive modulus

Prior to mechanical testing, cartilage and skin scaffolds were hydrated in PBS for 48 hours. Unconfined compression testing was performed using a rheometer (MCR 702, Anton Paar), compressing hydrated scaffolds to 40% of their original height at a quasi-static strain rate of 0.1% strain per second. Stress-strain data were recorded, and the compressive modulus was calculated from the linear region corresponding to 20-30% strain.

### 2.12 Shelf life

Cartilage and skin particle-only gECM wafers were fabricated and stored under dry conditions at room temperature. Mechanical compression testing, volume fraction analysis, and swelling characterization were performed immediately after fabrication (t0), as well as at 1 month and 3 months post-fabrication to evaluate scaffold property stability over time.

### 2.13 Cellular viability

Bovine chondrocytes and 3T3 mouse dermal fibroblasts were seeded onto cartilage and skin particle-only wafers, respectively, at a density of 4,000 cells/mm² based on wafer surface area. Following seeding, wafers were maintained under standard culture conditions (Chondrocytes: Dulbecco’s Modified Eagle DMEM-F12 supplemented with 10% FBS, 0.1% bovine serum albumin, 100 units/mL penicillin, 100 ug/mL strep, and 50 ug/mL ascorbate-2-phosphate; Dermal Fibroblast 3T3s: Dulbecco’s Modified Eagle DMEM-F12 supplemented with 10% FBS and 1% penicillin-streptomycin) for 14 days. Cell viability was assessed using LIVE/DEAD (Calcein AM, Ethidium Homodimer-1; ThermoFisher Invitrogen) staining at Days 3 and 14 post-seeding. Stained wafers were imaged using confocal microscopy (Nikon A1R, 10x objective, NA=0.45) to evaluate cell viability and spatial distribution within the wafer. Particles were visualized via autofluorescence using 405 nm excitation. Wafer surface area was measured from stereoscopic images acquired using a Leica microscope and quantified via thresholding in ImageJ. To evaluate changes in mechanical properties during culture, unconfined compression testing was performed (0.1% strain/sec, compressed to 40% strain) at each time point using a mechanical testing system (BOSE ElectroForce 5500). Compressive modulus was calculated between 20-30% compression.

### 2.14 Cartilage implantation application

A juvenile bovine knee joint was used to evaluate scaffold implantation. Articular cartilage on the femoral condyle was exposed, and two cylindrical defects were created using a biopsy punch. Hydrated and dry cartilage particle-only wafers were implanted into the defects. For the dry wafer, PBS was applied dropwise to induce *in situ* hydration following implantation.

### 2.15 Statistical analysis

RStudio was used to perform mixed-model analyses of variance (ANOVA) on linear models with treatment (fabrication type, timepoint, etc.) as the primary factor, and the resulting measurement variable (stiffness, volume fraction, swelling weight, etc.) as the response. Residuals were checked, and the data was transformed for non-normal distributed residuals. Tukey’s honestly significant difference (HSD) correction post hoc tests were performed. Significance was defined as p<0.05.

## 3. RESULTS

### 3.1 Decellularized cartilage and skin ECM yield size-controlled particles with tissue-specific swelling behavior

Decellularized extracellular matrix particles were generated from human skin and cartilage through viral inactivation, lyophilization, pulverization, and size-sorting to <250 µm (Figure 1A). Successful decellularization was confirmed by measuring residual double-stranded DNA content (Figure 1B). DNA levels in both tissues were below the accepted threshold of 50 ng/mg dry tissue [20], with cartilage containing 0.9 ng/mg and skin containing 8.8 ng/mg on average. Particle size distributions revealed significant swelling in both tissues (Figure 1C). Mean particle diameter increased by ∼50% in cartilage (173 µm to 261 µm) and ∼40% in skin (38 µm to 54 µm). Cartilage particles were consistently larger than skin particles in both non-swelled and swelled states. Together, these results demonstrate that decellularized ECM particles can be reproducibly generated with controlled size and tissue-specific swelling behavior, establishing them as reliable building blocks for forming cohesive particle-only wafers.

**Figure 1.**
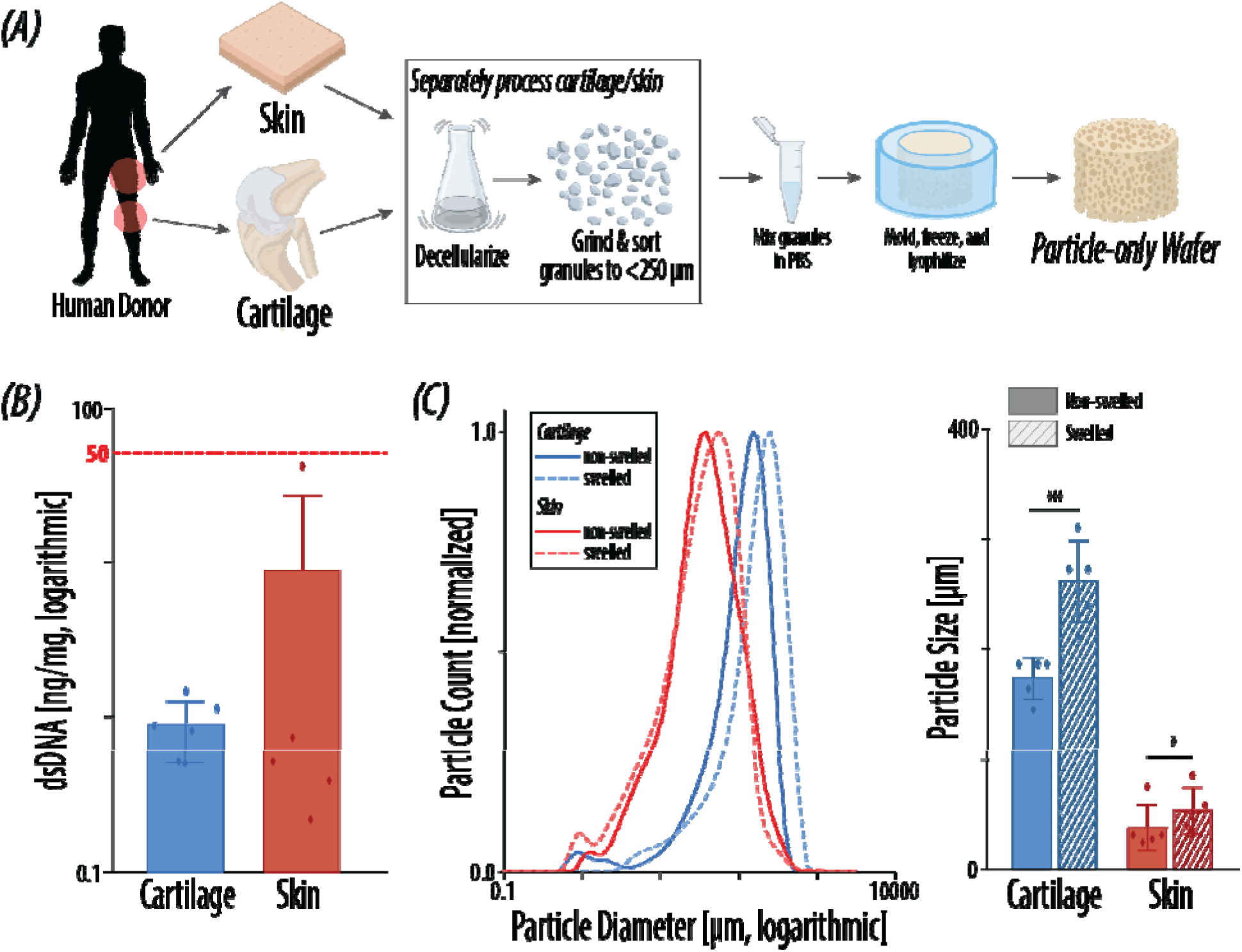
Tissue processing workflow and ECM particle characterization. (A) Tissues are decellularized and pulverized to particles <250 µm. ECM particles are suspended in PBS, then frozen and lyophilized to produce particle-only gECM wafers. (B) Double-stranded DNA content of decellularized tissues (N=5-6). (C) Particle size distributions and mean size of tissue particles in non-swelled and swelled states (N=5). Error bars=standard deviation, *p<0.05, **p<0.01, ***p<0.001.

### 3.2 Cartilage and skin ECM exhibit distinct, collagen-dominant proteomic profiles with tissue-specific matrisome composition

Proteomic analysis revealed distinct ECM composition between cartilage and skin tissues (Figure 2A). A heatmap of matrisome proteins showed clear differences in protein abundance by tissue type, reflecting divergent ECM profiles. The majority of detected proteins in both tissue were matrisomal, comprising 99.3% of total label free quantification (LFQ) intensity in cartilage and 99.7% in skin, while cellular proteins represented only a minor fraction, further supporting successful particle decellularization.

**Figure 2.**
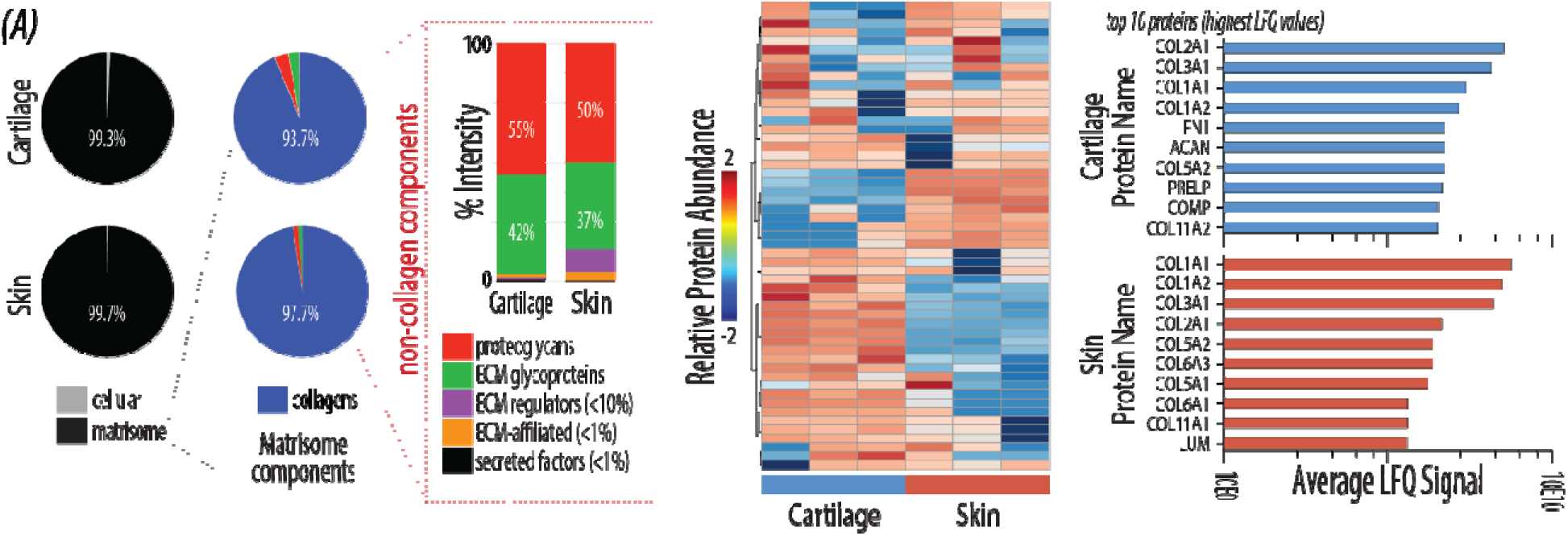
Proteomic characterization of cartilage and skin extracellular matrix. (A) Pie charts display average composition of each tissue type by LFQ signal: cellular vs. matrisome proteins and composition by matrisome subcategory. Composition of non-collagen matrsiome components is shown as percentage of total-non collagen matrisom protein LFQ intensity. Heat map showing the relative abundance of detected matrisome proteins in deceullularized cartilage and skin tissues. The top ten most abundant matrisome proteins in cartilage and skin tissues are ranked by average LFQ intensity (N=3).

Within the matrisome, collagens were the dominant component in both tissues. Among non-collagenous components, proteoglycans and ECM glycoproteins represented the largest fractions. Proteoglycans accounted for 55% of LFQ intensity for non-collagen ECM components in cartilage and 50% in skin, while ECM glycoproteins comprised 42% and 37%, respectively. The most abundant proteins differed between tissues. Cartilage ECM was enriched in collagen II (COL2A1, the primary load-bearing collagen in cartilage), fibronectin (FN1), and aggrecan (ACAN), whereas skin ECM was dominated by collagen I (COL1A1, COL1A2) and collagen III (COL3A1), which contribute to tensile strength in fibrous connective tissues [26]. These results confirm tissue-specific ECM composition despite both tissues being collagen-rich, and match the known composition of native cartilage and skin [27–29]. Overall, these findings confirm that decellularized ECM particles retain a highly ECM-rich and tissue-specific protein composition, preserving key biochemical cues that are expected to influence scaffold mechanics and cell-matrix interactions.

### 3.3 Cartilage and skin ECM particles display distinct microstructures and surface roughness that are preserved upon hydration

Scanning electron microscopy (SEM) revealed distinct microstructures between cartilage and skin ECM particles (Figure 3A). In the dry state, particle-only cartilage wafers exhibited a porous structure and particles surfaces appeared rough. Upon hydration, particles became more compact with smoother surfaces and wafer porosity was reduced. In contrast, skin particles displayed a dense, fibrous network that was largely preserved after hydration, and similarly to cartilage, surface features appeared smoother.

**Figure 3.**
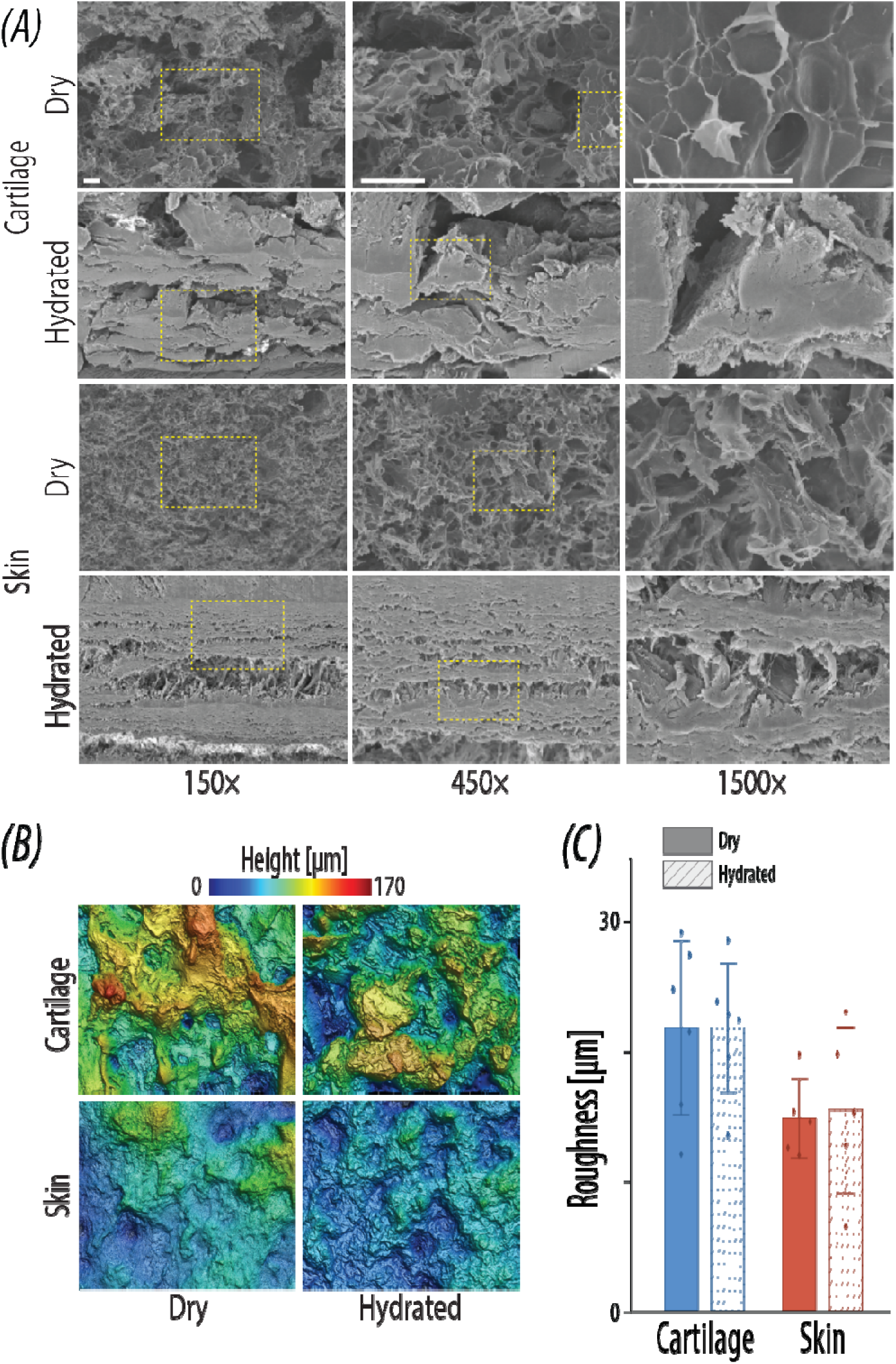
Microstructure and surface topology of cartilage and skin particle-only gECM wafers. (A) Scanning electron microscopy (SEM) images with increasing magnifications (150X, 450X, and 1500X) of dry and hydrated cartilage and skin particle-only gECM wafers. Scale bars=50 µm. (B) Three-dimensional surface topology maps showing surface roughness features of cartilage and skin particle-only wafers. (C) Quantification of surface roughness (Sa) for cartilage and skin gECM in dry and hydrated states (N=5-6). Error bars=standard deviation; *p<0.05, **p<0.01, ***p<0.001.

Surface topology analysis demonstrated heterogeneous roughness across both tissue types (Figure 3B). Cartilage particles exhibited higher surface roughness (Sa = 28.1 µm) compared to skin particles (Sa = 16.2 µm) (Figure 3C). Hydration did not significantly alter surface roughness in either tissue. Together, these results demonstrate that ECM particle microstructure and surface topology are tissue-specific and largely preserved upon hydration, indicating that particle-only wafers retain structural features relevant to cell-material interactions. These properties may influence cell adhesion, migration, and infiltration within the scaffold, ultimately impacting tissue integration and remodeling.

### 3.4 Scaffold fabrication method modulates swelling, structure, and mechanical properties in a tissue-dependent manner

Scaffold properties were strongly influenced by fabrication method, with distinct differences in swelling behavior, structure, and mechanical performance observed across scaffold types (Figure 4). All scaffolds exhibited significant increases in mean weight and volume within the first hour of hydration (Figure 4C). Cartilage scaffolds reached an equilibrium state after one hour, whereas skin scaffolds continued to swell over one week, particularly in wafer-based formats. The gECM hydrogel exhibited the greatest increase in weight (5.97x for cartilage; 6.8x for skin), while the gECM wafer exhibited the largest volume expansion (1.76x for cartilage; 1.62x for skin) (Table 1). Despite these differences, swelling followed consistent, predictable trends across fabrication methods, suggesting that scaffold behavior can be standardized for translational use.

**Figure 4.**
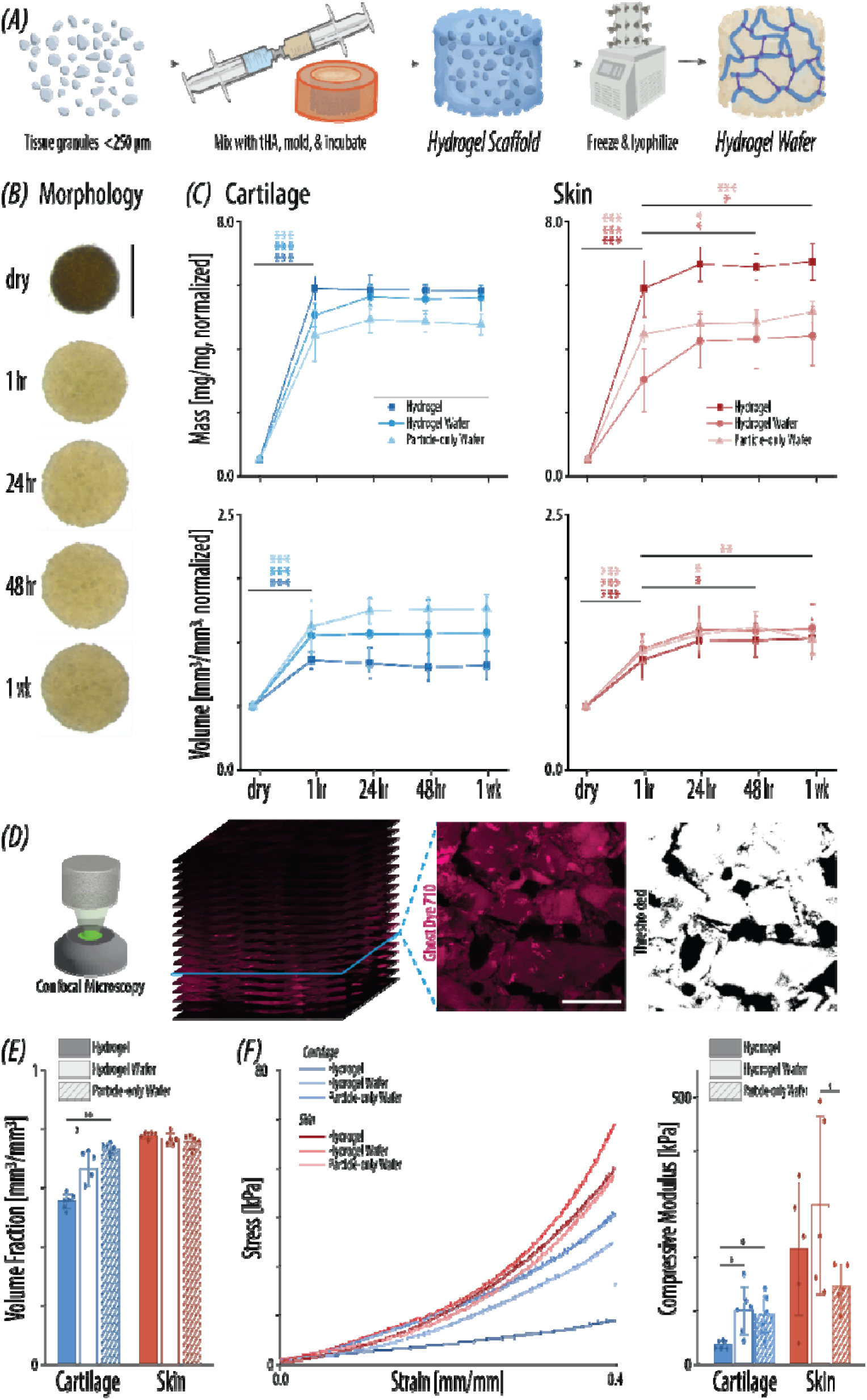
Comparison of gECM scaffold fabrication methods. (A) ECM particles are mixed with thiolated hyaluronic acid (tHA) and incubated at 37°C to produce crosslinked, gECM hydrogel scaffolds, which are then frozen and lyophilized to produce gECM hydrogel wafers. (B) Morphology of a cartilage particle-only gECM wafer during hydration over one week. Scale bar=5mm. (C) Normalized weight and volume of cartilage and skin ECM scaffolds measured over time during hydration for gECM hydrogel, gECM hydrogel wafer, and particle-only gECM wafer fabrication methods. (D) Z-stack, raw, and thresholded slice of scaffold structure. Images taken using confocal microscopy and thresholded using MATLAB to quantify volume fraction within a 15 µm depth. Scale bar=200 µm. (E) Volume fraction for cartilage and skin scaffolds of different fabrication methods. (F) Representative stress-strain curves and calculated compressive modulus (20-30% strain) for cartilage and skin scaffolds of different fabrication methods. Data for cartilage hydrogel and hydrogel wafer groups in panels C, E, and F were previously reported in Heye et al. [30] and are included here for comparison with newly generated cartilage particle-only gECM wafer data. All methods N=5-6; error bars=standard deviation; *p<0.05, **p<0.01, ***p<0.001.

**Table 1.** Mean change in scaffold weight and volume compared to dry (normalized) at 1 week

|  | gECM hydrogel | gECM hydrogel wafer | Particle-only gECM wafer |
| --- | --- | --- | --- |
| Cartilage weight | 5.97 ± 0.18* | 5.77 ± 0.37* | 4.98 ± 0.24 |
| Cartilage volume | 1.31 ± 0.11* | 1.57 ± 0.26* | 1.76 ± 0.07 |
| Skin weight | 6.80 ± 0.54 | 4.64 ± 0.74 | 5.34 ± 0.22 |
| Skin volume | 1.52 ± 0.13 | 1.59 ± 0.12 | 1.62 ± 0.09 |

\*Data for the cartilage gECM hydrogel and gECM hydrogel wafer groups were previously published in Heye et al. [30] and are reproduced here for comparison with newly generated cartilage particle-only gECM wafer data.

Biophysical and mechanical analysis revealed differences in particle volume fraction (Figure 4E) and bulk stiffness (Figure 4F) across fabrication methods. For cartilage, there were significant differences between the hydrogel scaffold and wafer-based formats. gECM wafers exhibited the highest volume fraction of particles-to-total-area (73.3%), followed by gECM hydrogel wafers (66.5%) and gECM hydrogels (55.3%). In contrast, skin scaffolds exhibited similar volume fractions across all fabrication methods, between 75-78%. Mechanical testing demonstrated that stiffness varied with both tissue type and fabrication method. Skin scaffolds were consistently stiffer than cartilage scaffolds, and wafer-based scaffolds exhibited greater stiffness than hydrogels. Among both tissues, gECM hydrogel wafers exhibited the highest stiffness (99.8 kPa for cartilage; 296.2 kPa for skin). Together, these results demonstrate that fabrication method modulates scaffold structure and mechanics, with particle-dense wafer formats promoting increased structural organization and stiffness. Together, these results demonstrate that fabrication method governs scaffold structure and mechanics, with particle-dense wafer formats promoting increased organization and stiffness. Importantly, these findings show that particle-only scaffolds can achieve mechanically relevant performance without polymer reinforcement, supporting their viability as a simplified alternative to hydrogel-based systems.

### 3.5 Particle-only gECM wafers maintain swelling behavior, structure, and mechanical properties over 3 months of storage

Wafer properties remained stable over a 3-month storage period, with minimal changes in swelling behavior, structure, and mechanical properties (Figure 5). Swelling behavior was consistent across timepoints, with both cartilage and skin wafers exhibiting a significant increase in mean weight and volume within 30 minutes of hydration, followed by gradual increases over one week. Despite this rapid initial increase, representative images revealed that wafers were not fully swelled at 30 minutes, but reached a fully swelled state by 8 hours, consistent with the plateau observed in quantitative measurements (Figure 5A). Overall swelling remained similar over 3 months, with cartilage particle-only wafers increasing to ∼5.2x dry weight and ∼1.6x dry volume, and skin to ∼5.1x weight and ∼1.6x volume (Table 2).

**Figure 5.**
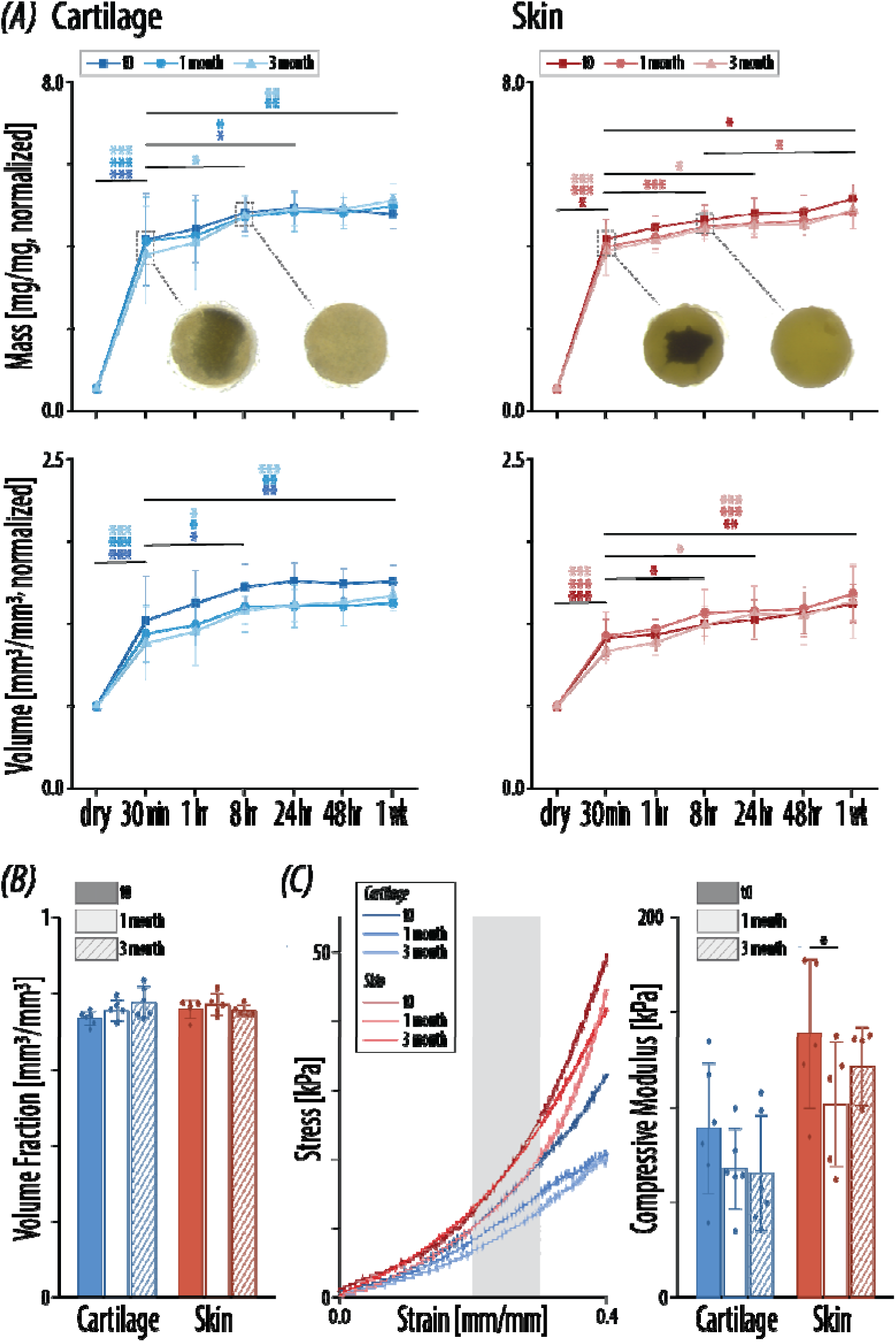
Shelf-life stability of cartilage and skin particle-only gECM wafers. (A) Normalized weight and volume of cartilage and skin ECM wafers measured over time during hydration at t0, 1 month, and 3 months. Representative images of wafers are shown at 30-minute and 8-hour timepoints. (B) Wafer volume fraction quantified from confocal microscopy imaging and MATLAB thresholding for cartilage and skin particle-only wafers at t0, 1 month, and 3 month timepoints. (C) Representative stress-strain curves and calculated compressive modulus (20–30% strain) for cartilage and skin particle-only at t0, 1 month, and 3 months. All methods N=5-6; error bars=standard deviation; *p<0.05, **p<0.01, ***p<0.001.

**Table 2.** Mean change in gECM particle-only wafer weight and volume compared to dry (normalized) at 1 week.

|  | t0 | 1 month | 3 months |
| --- | --- | --- | --- |
| Cartilage weight | $4.98 \pm 0.24$ | $5.17 \pm 0.22$ | $5.30 \pm 0.26$ |
| Cartilage volume | $1.76 \pm 0.07$ | $1.62 \pm 0.09$ | $1.67 \pm 0.07$ |
| Skin weight | $5.34 \pm 0.22$ | $5.03 \pm 0.29$ | $5.08 \pm 0.29$ |
| Skin volume | $1.62 \pm 0.09$ | $1.68 \pm 0.12$ | $1.64 \pm 0.15$ |

Structural stability was maintained over time, with no significant changes in volume fraction for either tissue (Figure 5B). Cartilage wafers exhibited a slight increase in volume fraction from 73.3% at t0 to 77.5% at 3 months, while skin wafers remained relatively constant around 76% on average. Mechanical properties decreased with time (Figure 5C). Cartilage wafer stiffness decreased from 93.4 kPa at t0 to 68.5 kPa at 3 months, while skin decreased from 145.5 kPa to 127.6 kPa. These results suggest that particle-only gECM wafers maintain structural integrity and decrease slightly in strength over time, supporting their potential for off-the-shelf use and translational applications.

### 3.6 Particle-only gECM wafers support cell viability

Particle-only gECM wafers supported cell viability over 14 days in culture (Figure 6A–B). LIVE/DEAD staining showed predominantly viable cells at both Day 3 and Day 14 in cartilage and skin wafers. Chondrocytes were distributed throughout cartilage wafers, including both surface and interior regions. In contrast, fibroblasts on skin wafers were primarily localized to the surface, with more limited penetration into the interior. This difference may be attributed to variations in particle size and volume fraction between tissues, as larger, irregularly shaped cartilage particles create greater interparticle space that facilitates cell infiltration, whereas the denser, more entangled structure of skin wafers limits penetration.

**Figure 6.**
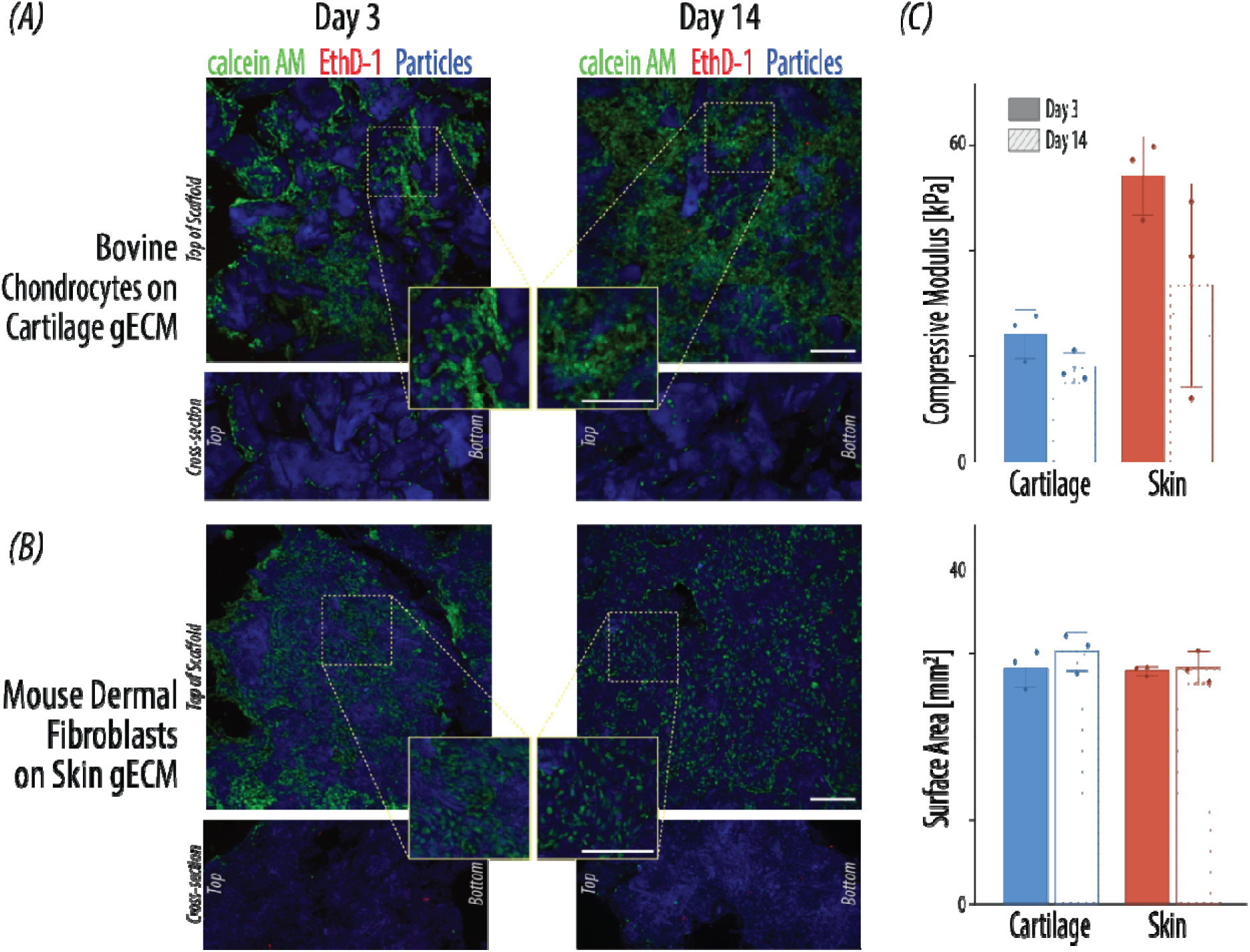
Cytocompatibility of cartilage and skin particle-only gECM wafers. (A) LIVE/DEAD fluorescence imaging of bovine chondrocytes cultured on cartilage particle-only wafers and (B) 3T3 mouse dermal fibroblasts cultured on skin particle-only wafers at Day 3 and Day 14. Live cells are labeled with calcein AM (green), dead cells with ethidium homodimer-1 (red), and particles are visualized via autofluorescence using 405 nm excitation (blue). Representative images show wafer top views and cross-sections. Scale bars=200 µm. (C) Compressive modulus of cartilage and skin particle-only wafers measured at Day 3 and Day 14 during cell culture. (D) Quantification of wafer surface area for cartilage and skin particle-only wafers over the culture period. Cell N=1; Tissue N=3; Error bars=standard deviation.

Particle-only wafer mechanical properties decreased over time in culture (Figure 6C). Cartilage wafers stiffness decreased from 23.6 kPa at Day 3 to 17.5 kPa at Day 14, while skin decreased from 52.9 kPa to 32.6 kPa. Surface remained stable over time for both tissue types (Figure 6D). Changes in stiffness may result from relaxation of particles within the wafer over culture time. Overall, these results demonstrate that particle-only gECM wafers provide a biologically supportive environment suitable for tissue engineering applications.

### 3.7 Particle-only gECM wafers conform to *ex vivo* cartilage defect geometry and remain cohesive under implantation conditions

Particle-only gECM wafers successfully conformed to defect geometry and remained localized within cartilage defect sites under both dry and pre-hydrated conditions (Figure 7). Dry wafers expanded *in situ* following PBS hydration, while pre-hydrated wafers were easily positioned (press-fit) within the defect. In both cases, the wafers conformed to defect geometry and provided smooth topological surfaces comparable to surrounding native cartilage. These results demonstrate that particle-only gECM wafers can be readily handled, conform to defect geometry, and remain cohesive under implantation conditions, supporting their feasibility for practical use in tissue repair settings.

**Figure 7.**
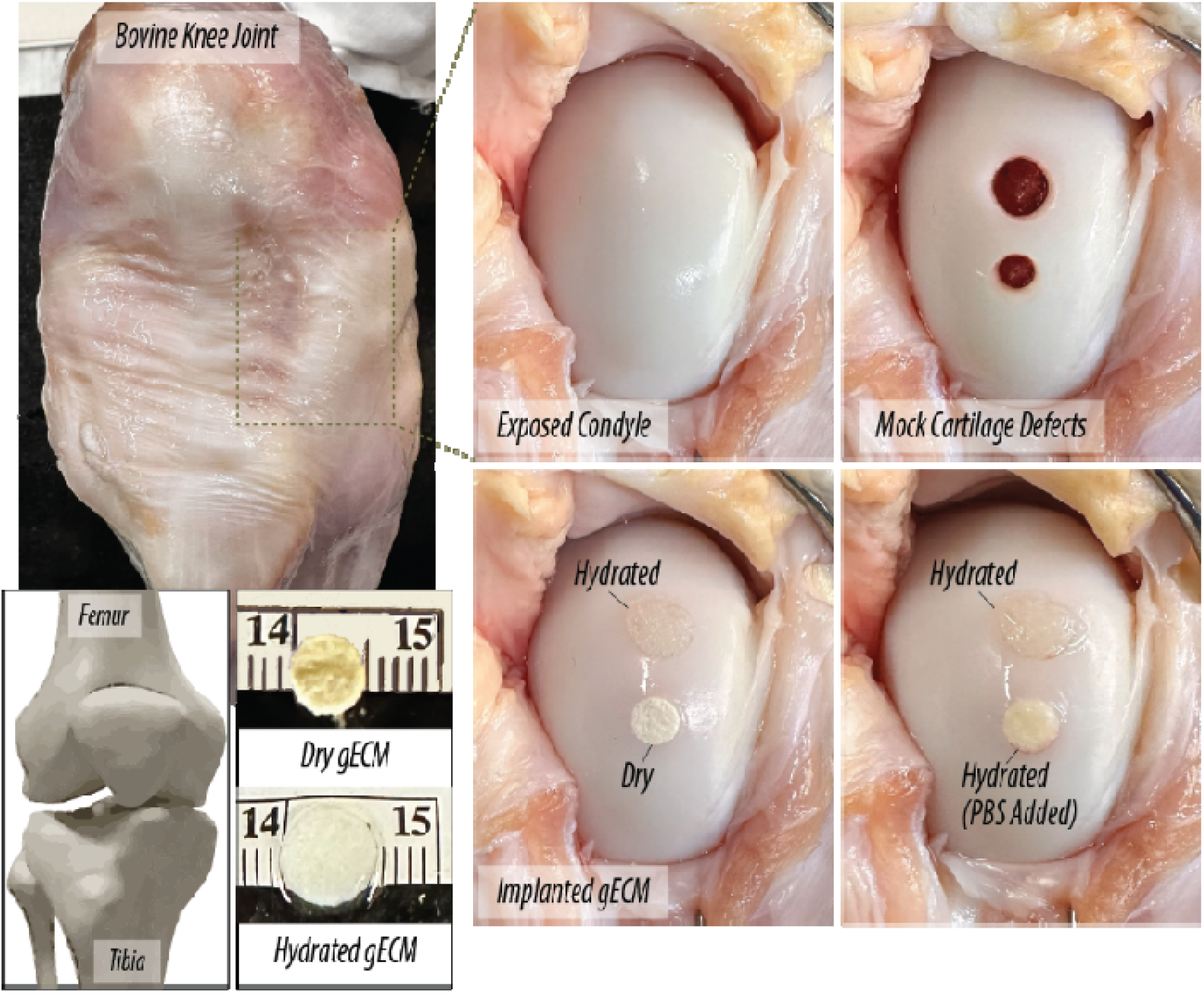
Demonstration of particle-only gECM wafer implantation in a bovine cartilage defect model. Implantation of cartilage particle-only wafers in a bovine knee joint. The femoral condyle was exposed and mock cartilage defects were created within the articular surface. Both dry and hydrated particle-only wafers were placed within the defects. Images show the exposed condyle, defect creation, and placement of wafers in dry and hydrated states.

## 4. DISCUSSION

This study evaluated whether particle-only gECM wafers could provide a simplified alternative to hydrogel-based gECM biomaterials while maintaining structural, mechanical, and biological performance. Particle-only gECM wafers were successfully fabricated from cartilage and skin ECM particles, forming cohesive structures without a secondary polymer network. These wafers exhibited tissue-dependent swelling and mechanical behavior, maintained structural properties over 3 months of storage, supported cytocompatibility, and remained cohesive during implantation. Together, these findings demonstrate that particle-only gECM wafers function as stable, ECM-rich biomaterials with strong translational potential.

Scaffold properties were strongly influenced by fabrication method, highlighting the role of processing in determining material behavior. The greater swelling and lower stiffness observed in gECM hydrogels suggest that non-lyophilized gECM polymer networks are less structurally constrained than densely packed wafer-based systems. This is likely due to ice crystal templating and dehydration-induced matrix compaction [16,31] in the freeze-drying step, which promotes structural densification and enhances load-bearing capacity relative to hydrated polymer networks. Notably, gECM hydrogel wafers exhibited the highest stiffness, indicating that combined polymer crosslinking and lyophilization may produce synergistic effects. In past work, we found that higher particle volume fraction is correlated with higher bulk stiffness through percolation [32]. This may explain some differences observed in both volume fraction and stiffness across fabrication methods, especially in cartilage scaffolds. Importantly, previous work found that once a critical particle volume fraction is reached, a continuous, load-bearing network can form through particle-particle contacts alone. Under these conditions, mechanical integrity is governed primarily by particle packing rather than the presence of a secondary polymer network [32].This may explain the minimal differences observed between hydrogel-based and particle-only wafers once this threshold is exceeded [33]. These results further support that a continuous polymer phase is not required for scaffold cohesion or mechanical integrity.

While gECM hydrogels are stabilized through thiol crosslinking [13], the formation of cohesive particle-only wafers is likely governed by a combination of physical packing and compositional mechanisms. During particle-only wafer fabrication, particles are confined within molds at relatively high mass-to-volume ratios. Under high-pressure lyophilization, particle-only gECM wafers likely experience further particle compaction leading to jamming, or dynamic arrest where particle motion is constrained by neighboring contacts [14]. Under these conditions, mechanical behavior is governed by normal and tangential forces at particle interfaces, including frictional contacts and particle deformation, which together enable formation of a continuous, load-bearing network without the need for chemical crosslinking [15,34,35]. gECM hydrogel wafers are likely stabilized through similar particle jamming in addition to hydrogel crosslinking and densification. Freeze-drying densifies the hydrogel component through ice crystal formation and sublimation, which may facilitate physical bonding and stabilize particle assemblies, as observed in freeze-dried collagen and ECM-derived sponge scaffolds [16–18]. This framework is consistent with the high volume fractions and mechanical stability observed in gECM wafers compared to non-lyophilized gECM hydrogel scaffolds, supporting the role of particle jamming in governing cohesion.

In addition to these physical packing mechanisms, ECM composition may further reinforce wafer stability. ECM-derived particles retain native proteins, proteoglycans, and structural features that enable biologically relevant interparticle interactions, including collagen fiber entanglement and surface adhesion [18,36]. These interactions may be particularly important in skin-derived scaffolds, where collagen I fibers promote entanglement, whereas cartilage scaffolds, composed of collagen II within a proteoglycan-rich matrix, may rely more heavily on particle packing and surface interactions. This interpretation is consistent with the observed tissue-dependent differences in scaffold structure and mechanical behavior. While the volume fraction is significantly different between gECM hydrogel and wafer formats in cartilage, it remains constant across fabrication types for skin. Together, these findings suggest that wafer cohesion arises from the combined effects of particle jamming, processing-induced densification, and ECM-specific interparticle interactions.

These proposed cohesion mechanisms are further supported by tissue-specific microstructural and differences observed by SEM. Cartilage particle-only wafers exhibited a more porous architecture with irregular, rough particle surfaces, whereas skin particle-only wafers displayed denser and more fibrous morphology. The rougher cartilage particle morphology may increase frictional contacts and mechanical interlocking between neighboring particles, while the fibrous structure and smaller particle size of skin may promote particle entanglement and increase particle-particle contact throughout the scaffold. These structural features were largely preserved following hydration, suggesting that particle-scale interactions remain relevant under physiological conditions. These observations align with prior work showing that particle size, morphology, and packing behavior influence mechanical stability and transport in porous biomaterials [37], highlighting the combined roles of ECM composition and particle architecture in governing wafer performance.

Shelf-life results provide additional insight into the stability and evolution of particle-only wafers over time. Swelling behavior and volume fraction remained stable over 3 months, indicating that the overall wafer architecture was preserved during storage. Notably, although wafers exhibited rapid increases in weight and volume within the first 30 minutes of hydration, representative imaging demonstrated that full swelling was not achieved until approximately 8 hours, suggesting that early hydration does not reflect a fully equilibrated wafer structure. This distinction may be important for both *in vitro* preparation and potential clinical use, where complete equilibration could influence handling and implantation timing. Wafer stiffness decreased modestly in both tissues, which may reflect microstructural relaxation or changes in particle contacts. Importantly, wafer cohesion and function were preserved during dry storage at room temperature. Lyophilized biomaterials, including collagen-based sponges and ECM-derived scaffolds, have been widely reported to maintain structural and functional properties over extended storage periods [37–39]. The ability to store gECM wafers in a dry state and rehydrate them without loss of structure represents a significant translational advantage, particularly for applications requiring off-the-shelf availability.

Building on their structural stability, particle-only gECM wafers supported cell viability over 14 days, indicating that the fabrication and processing methods did not introduce cytotoxic effects. However, cell distribution differed between tissues, with chondrocytes infiltrating cartilage wafers while fibroblasts remained primarily localized to the surface of skin wafers. These differences likely arise from microstructural factors such as particle size and packing density, which govern pore architecture and interparticle spacing. Smaller, elongated skin particles promote denser packing and reduced void space, limiting infiltration, whereas larger cartilage particles create greater interparticle spacing that facilitates cell penetration. Scaffold pore size and interconnectivity are well-established regulators of cell infiltration and tissue integration, with smaller pores and higher packing densities limiting cellular migration [40,41]. Processing parameters such as mass-to-volume ratio and freezing conditions may further influence pore size and connectivity and therefore represent potential design variables for tuning cellular access to the wafer interior. Conventional 2D culture systems are widely used due to their simplicity but lack physiological relevance, often resulting in altered cell morphology and loss of tissue-specific function over time, such as chondrocyte dedifferentiation [42]. In contrast, 3D encapsulation better supports native cell behavior by providing structural and biochemical cues but typically requires more complex fabrication which limits accessibility [43,44]. Particle-only wafers represent potential for an intermediate 2.5D environment, in which cells are seeded on the surface while still interacting with a porous, structural, and ECM-rich substrate. This configuration enables improved biological relevance compared to 2D systems while maintaining accessibility for imaging and analysis, and may explain the surface-localized yet viable cell behavior observed in skin wafers.

These findings highlight how wafer architecture can be leveraged to direct cell-material interactions, enabling design of systems that support either surface-guided growth or deeper tissue integration. Wafer stiffness decreased over time in culture, likely reflecting matrix remodeling, progressive particle relaxation during hydration, or early-stage degradation. Changes in matrix stiffness are known to influence cell behavior and may arise from both material degradation and cell-mediated remodeling processes [45,46]. While such softening is expected for biologically derived materials, its implications for long-term mechanical function, particularly in load-bearing applications, require further investigation. The implantation study demonstrated that particle-only wafers are mechanically robust and conformable during delivery to an *ex vivo* joint, supporting their potential for practical *in vivo* use. Wafers maintained structural integrity during handling and implantation and could be delivered either in a hydrated state or as dry constructs that expand *in situ*. These characteristics suggest that particle-only gECM wafers may be suitable for minimally invasive or press-fit applications and reinforce their potential as off-the-shelf biomaterials.

In addition to demonstrated functional and application-relevant properties, these observed structure-function relationships suggest that processing parameters may represent key opportunities to tune wafer properties. For example, the mass-to-volume ratio used during fabrication likely influences particle packing density, volume fraction, and stiffness, as suggested by preliminary observations. Similarly, freezing conditions may affect pore formation and wafer architecture. Isotropic freezing may produce more uniform pore structures, whereas anisotropic freezing could introduce directional porosity and alter volume fraction [31]. While not systematically explored here, these parameters represent important design variables for tuning wafer structure and mechanical performance in future work.

Overall, particle-only gECM wafers eliminate the need for synthetic polymers and crosslinking chemistries while retaining high ECM content and functional performance. By avoiding polymer carriers, these systems may preserve direct cell-matrix interactions and reduce the risk of altering ECM composition or introducing cytotoxic byproducts associated with chemical modification [47]. This reduction in material complexity may also simplify manufacturing and regulatory pathways while maintaining biologically relevant matrix cues. In addition, the ability to fabricate and store these wafers in a lyophilized state enables room-temperature storage and off-the-shelf availability, addressing a key limitation of hydrogel-based systems that require hydrated conditions and more complex handling. Furthermore, initial observations suggest that this platform may be extended to additional tissue types, such as kidney, liver, and bone. However, several limitations should be considered. This study focused on two tissue types and short-term *in vitro* evaluation, and mechanical testing was limited to compression whereas tension may be more applicable for elastic tissues like skin. In addition, donor samples were predominantly male, and future work should incorporate a more balanced representation of biological sex, as sex-specific differences in ECM composition, structure, and remodeling behavior may influence wafer properties and cell-matrix interactions. Limited cell infiltration in skin wafers suggests that further optimization of wafer architecture may be required. Furthermore, while cytocompatibility was demonstrated, functional cell behavior, including gene expression and matrix degradation and deposition, was not assessed and will be important to evaluate in future studies. Future work should focus on expanding to additional tissue types, evaluating long-term functional performance, and systematically tuning wafer architecture through control of particle size, packing density, and processing conditions.

## 5. CONCLUSION

Particle-only gECM wafers were developed as cohesive, ECM-rich biomaterials without secondary polymer networks. Decellularized ECM particles retained tissue-specific composition and structure, providing a biologically relevant foundation for scaffold formation. Through freeze-drying and high-density packing, these wafers formed stable, load-bearing constructs, and demonstrate that a continuous polymer network is not required for cohesion, mechanical integrity, or biological support. The wafers exhibited distinct structural and mechanical properties, maintained stability over extended room temperature storage, and supported cell viability and dynamic cell-material interactions *in vitro*. Differences between cartilage and skin wafers highlight the role of tissue-specific ECM composition and architecture in governing wafer behavior, including particle interactions, cell infiltration, and mechanical performance. Importantly, particle-only gECM wafers conformed to defect geometry and remained cohesive under physiologically relevant conditions, supporting their feasibility for tissue engineering applications. By eliminating the need for synthetic polymers and chemical crosslinking, this platform offers a simplified and more translationally favorable approach to ECM-based biomaterials. Overall, this work establishes particle-only gECM wafers as a structurally and biologically functional platform for tissue engineering and a foundation for future gECM-based material development.

## Acknowledgments

Donor tissues provided by AlloSource are gratefully acknowledged. Mass Spectrometry was performed by the Mass Spectrometry Proteomics Shared Resource Facility [RRID SCR_021988] at the University of Colorado Anschutz.

## Funding

The authors gratefully acknowledge funding from the following sources: National Institutes of Health R01 AR083379 and U01 AR082845 (CPN), National Science Foundation CMMI 2212121 (CPN), and National Science Foundation Graduate Research Fellowship (JOH).

## Author Contributions

Conceptualization: SAB, CPN

Methodology: SAB, JOH, SES, MCM, CPN

Visualization: SAB, CPN

Funding acquisition: CPN

Supervision: JOH, SES, CPN

Writing – original draft: SAB

Writing – review & editing: All authors

## Competing interests

Authors JOH and CPN have equity in TissueForm, Inc.

